# Pathogenic Keratinocyte States and Fibroblast Niches Define the Tissue Microenvironment in Severe Hidradenitis Suppurativa

**DOI:** 10.1101/2025.11.11.682326

**Authors:** Xinyi Du-Harpur, Clarisse Ganier, Pavel Mazin, Chris Cheshire, Jeyrroy Gabriel, Ellie Rashidghamat, Nicholas M Luscombe, Magnus Lynch, Fiona Watt

**Author notes:** These authors contributed equally to this work (Xinyi Du-Harpur, Clarisse Ganier). **Correspondence: Xinyi Du-Harpur,** St. John’s Institute of Dermatology, King’s College London, London, United Kingdom.; **Clarisse Ganier,** Meta-organism Unit, Immunology Department, Institut Pasteur, Paris, France.; and **Fiona M. Watt,** The Directors’ Research Unit, European Molecular Biology Laboratory, Meyerhofstr. 1, Heidelberg 69117, Germany.

## Abstract

Hidradenitis suppurativa (HS) is a chronic inflammatory skin disease characterized by recurrent painful abscesses and tunnels in flexural sites. The mechanisms driving HS pathogenesis, particularly the roles of keratinocytes and fibroblasts in the HS inflammatory microenvironment, remain poorly understood. To characterise the cellular and molecular landscape of HS, we analyzed lesional skin from severe HS patients using single-cell RNA-sequencing and spatial transcriptomics to identify keratinocyte states and patterns of cellular interactions, with a focus on fibroblast-keratinocyte crosstalk. We identified a migratory S100+ pathogenic keratinocyte state enriched in HS lesions and observed spatially distinct fibroblast populations associated with different tissue compartments. COL6A5+ fibroblasts were preferentially co-localised with undifferentiated keratinocyte states, whereas APOD+ fibroblasts with enriched in immune cell-rich regions containing activated B and plasma cells consistent with tertiary lymphoid organ (TLO)-like structures. Ligand-receptor inference and spatial co-localisation analyses predicted extensive fibroblast-keratinocyte and fibroblast-immune interactions within HS lesions. Together, these data define the cellular organisation of the HS tissue microenvironment and identify spatially distinct stromal and epithelial niches associated with severe disease. These findings provide a framework for understanding how epithelial, stromal and immune cell populations coexist within HS lesions and may help to explain the limited efficacy of therapies targeting individual inflammatory pathways in the treatment of HS.

## Introduction

Hidradenitis suppurativa (HS) is a chronic inflammatory skin disease that affects approximately 1% of individuals in western countries (Ingram, 2020). It is characterised by the formation of painful nodules, abscesses and tunnels in flexural sites such as axillae and groin, leading to considerable psychosocial morbidity and impact on daily activities (Nguyen et al., 2021). HS is also associated with significantly increased cardiovascular adverse events, increased cancer risk and reduced life expectancy (Tiri et al., 2019).

HS pathogenesis originates in the pilosebaceous unit, where altered epidermal keratinisation, dysregulated innate immunity and dysbiosis leads to comedogenesis and inflammation (Frew, 2020). Severe HS (Hurley stage III) is characterised by the formation of tunnels, whereby keratinocytes invade the dermis and form structures that generate inflammatory exudate and propagate inflammation (Navrazhina et al., 2021). Dysbiosis, specifically an increased abundance of rare anaerobes such as *Porphyromonas* or *Prevotella*, is a feature of HS, with perilesional tissue demonstrating microbiome shifts reminiscent of lesional tissue (Agbogan et al., 2026; Riverain-Gillet et al., 2020).

Increased inflammation from immune signaling is observed in HS, with recent transcriptomic studies focusing on the role of polarised macrophages, B cells and tertiary lymphoid organs (TLOs) (Gudjonsson et al., 2020; Lowe et al., 2024; Mariottoni et al., 2021; Yu et al., 2024), which represent ectopic germinal center-like aggregates of immune cells and can arise in the context of chronic inflammation (Dong et al., 2023). Recently approved therapeutic options for severe HS are based on inhibition of IL-17 and TNF-alpha, and there are also promising clinical investigations into targeting JAK/STAT signalling (Krueger et al., 2024). However, these treatment options are often only partially effective.

Over the past two decades, studies characterising the genetic architecture of HS have repeatedly identified epidermal differentiation as a key pathogenic factor in HS, highlighting in particular the important role of γ-secretase in disease. γ-secretase in an intramembranous protease complex, which can cleave numerous transmembrane proteins including Notch receptors and cadherin (Pink et al., 2013). In mouse models, alterations in γ-secretase function lead to follicular keratinisation, epidermal cysts, epidermal hyperplasia and lack of sebaceous glands (Pan et al., 2004). A recent genome-wide association meta-analysis has confirmed the importance of both Notch and Wnt/β-catenin signaling pathways as being key to HS pathogenesis (Kjærsgaard Andersen et al., 2025). A previous GWAS identified SOX9 and KLF5 as putative causal genes in HS (Sun et al., 2023). SOX9 is a pioneer transcription factor that regulates fate determination of hair follicle outer root sheath keratinocytes (Yang et al., 2023). Hyperplasia of the outer root sheath is the characteristic histological feature of the very earliest stages of HS (Dunstan et al., 2021), and is observed clinically as comedone formation.

In light with those findings, we used single-cell RNA sequencing (scRNAseq) and spatial transcriptomics (ST) to characterise the cellular landscape of severe HS, with a focus on epithelial differentiation and stromal-epithelial interactions. We identified keratinocyte states with transcriptional features consistent with pathogenic behaviour in HS lesions, and examined how distinct fibroblast populations are spatially associated with epithelial and immune compartments. These findings support a model in which multiple interacting epithelial, stromal and immune compartments contribute to HS pathology, and may help to explain the limited efficacy of therapeutic strategies targeting individual inflammatory pathways in isolation, which contributes to the substantial healthcare burden associated with HS (Garg et al., 2020).

## Results

### Characterisation of cellular heterogeneity in lesional HS tissue

ScRNAseq (3’) data was generated from lesional axillary skin of three donors with severe HS (Figure 1a, Supplementary Figure 1a-b). Following quality control, dimensionality reduction and clustering in Seurat, cell types were annotated based on analysis of differentially expressed genes and canonical marker expression. This identified seven keratinocyte clusters, three fibroblast clusters as well as clusters corresponding to melanocytes, pericytes, endothelial cells (vascular and lymphatic) and multiple immune populations including Langerhans cells, macrophages, mast cells, B and plasma cells (Figure 1b-c).

**Figure 1.**
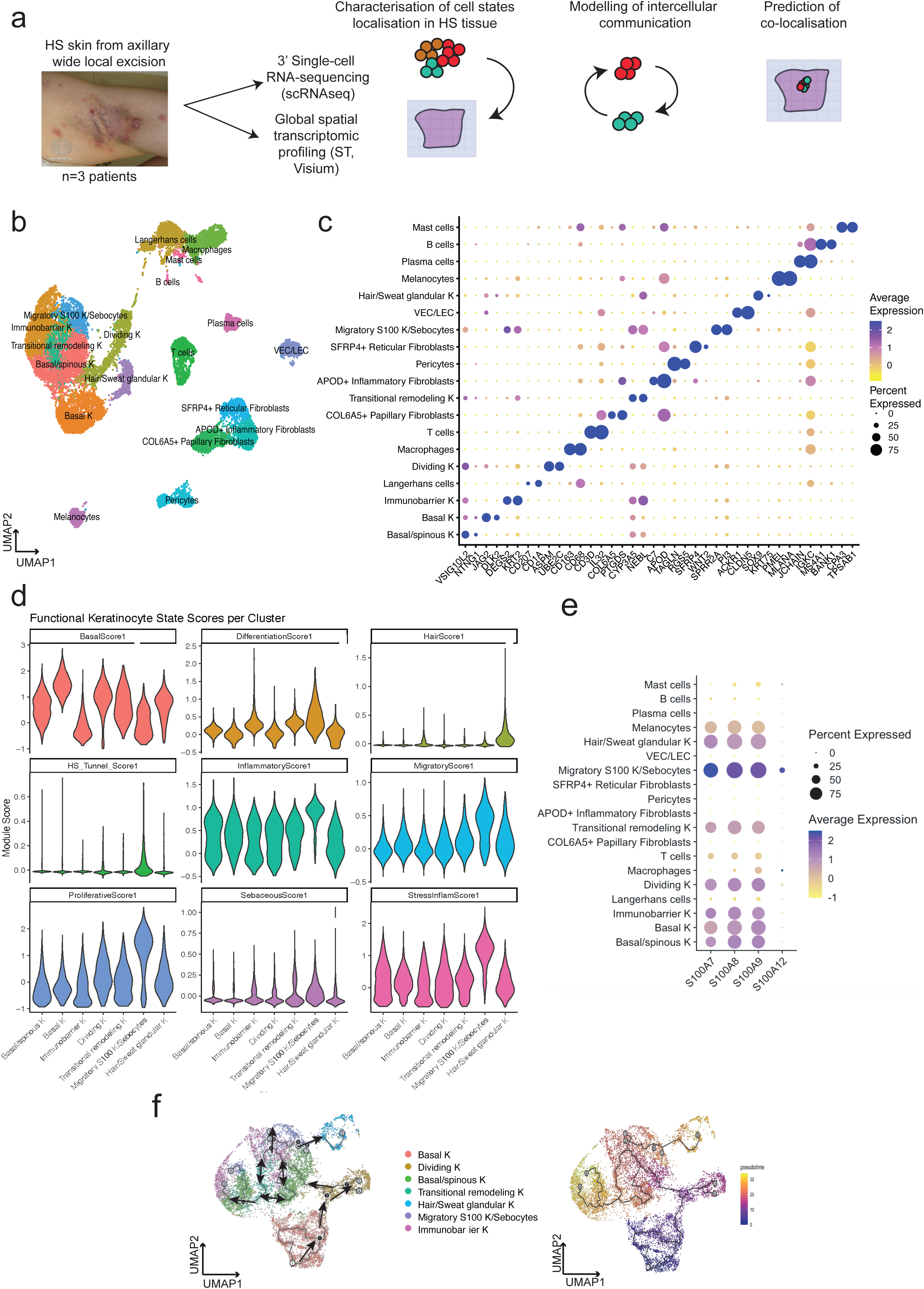
Single-cell landscape of HS skin reveals a distinct HS-associated pathogenic keratinocyte state. **(a)** Skin samples were obtained from 3 donors with severe HS undergoing axillary wide local excision surgery. ScR-NAseq (3’) and spatial transcriptomics (Visium) were performed on HS skin dissociated cells and 10um cryosections. Cell2location, cell-cell interaction and co-localisation analysis were undertaken to identify the location, identity and function of cells in HS skin. **(b)** Uniform Manifold Approximation and Projection (UMAP) and clustering of 20,003 cells from 3 donors representing 11 skin cell types and 19 cell populations. Each cell is represented by a single point within 2-dimensional space on the UMAP plot, with proximity of points plotted according to global transcriptional similarity. **(c)** Dot plot of established marker genes representative of each cell population. In this representation, the percentage of cells expressing the marker (diameter) and the average log2 normalized expression (color) are shown. **(d)** Violin plots demonstrating gene expression within 7 keratinocyte clusters, of signature functional gene modules (Table 1) related to keratinocyte state characteristics. These included functional modules representing proliferative, basal, differentiated, hair, sweat and sebaceous, as well as HS specific modules with tunnel, inflammatory, migratory and stress response genes. **(e)** Dot plot showing high expression of S100A7, S100A8, S100A9, and S100A12 in the pathogenic keratinocyte state of HS skin. **(f)** Single-cell trajectory gene analysis using Monocle 3 colored by keratinocyte state. Root node (white) indicates basal keratinocyte with lines indicating trajectory branch-es (arrows added for clarity) representing cellular decisions. Monocle pseudotime plot, where cells are colored according to differentiation state relative to the cell of origin.

### Transcriptional and spatial identification of a migratory S100+ Keratinocyte state in HS

Keratinocyte clusters were annotated according to their mean expression of gene signature modules representing functional characteristics (Table 1). The seven keratinocyte clusters identified were further characterized with proliferative, basal, differentiated, hair, sweat and sebaceous scores derived from these canonical marker gene modules (Figure 1d). HS associated gene signature modules included tunnel-associated, inflammatory, migratory and stress-response gene sets. We were also able to identify a keratinocyte state characterised by high expression of proliferation, differentiation, migration, inflammation and stress response genes including S100A7, S100A8, S100A9 and S100A12 (Figure 1e). This cell state simultaneously showed high expression of sebocyte and HS tunnel genes (Figure 1d) and was therefore annotated Migratory S100+ K/Sebocytes. This pattern of expression is consistent with recent molecular studies demonstrating glandular / appendageal gene expression patterns within HS tunnel epithelium (Lin et al., 2025). Similar S100-associated inflammatory keratinocyte populations arising from reprogrammed basal progenitor cells have been described by recent transcriptomic studies of HS lesions (L. Jin et al., 2023). Based on its transcriptional profile and enrichment within lesional tissue, we classified this cluster as a pathogenic keratinocyte state associated with HS.

Lineage trajectory and pseudotime analysis using Monocle (Figure 1f) suggested that basal keratinocytes transition through dividing, spinous and intermediate migratory keratinocyte (K) states prior to differentiating into three differentiated states: immunobarrier K, migratory S100+ K and hair / sweat glandular K. A transitional remodeling keratinocyte cluster was positioned within a closed loop trajectory, which reflect increased transcriptional plasticity within HS lesions. Similar epithelial plasticity and reprogramming of basal progenitor cells has been described in HS using single-cell and epigenetic analyses (L. Jin et al., 2023), and keratinocyte plasticity has also been demonstrated in models of epidermal injury and repair, including the recruitment of GATA6+ lineage keratinocytes from appendageal compartments during wound-induced remodeling (Bernabé-Rubio et al., 2023).

Having catalogued keratinocytes populations from HS lesional tissues, we localised them within the tissue sections using 10X Visium ST. To estimate the spatial distribution of the different cell types identified in our scRNAseq dataset, we used a cell2location algorithmic approach (Kleshchevnikov et al., 2022) to map the annotated scRNAseq clusters onto the Visium data. ST libraries were prepared from four HS skin samples, and QC metrics confirmed good library quality (Supplementary Figure 1c-d). Cell2location was then used to infer the cell composition of each Visium spot based on the scRNAseq reference, allowing spatial prediction of cell populations across the tissue sections.

Spatial prediction suggested that dividing K were sporadically distributed in the epidermis and within HS tunnels identified by H&E. Basal K and basal/spinous K were detected in the basal layer of both the interfollicular epidermis and predicted within HS tunnels. Transitional remodeling K were also predicted throughout the interfollicular epidermis layers except for the granular layer; with a similar distribution being predicted within HS tunnel epithelium. Immunobarrier K were enriched with the most differentiated and granular layer of the interfollicular epidermis, and the luminal layer of HS tunnels. Hair and sweat gland-associated K were predominantly localised to appendageal structures identified by histology of healthy samples (Supplementary Figure 2a), and were not strongly predicted within tunnel regions in the HS sections. In contrast, migratory S100+ K showed relative enrichment within specific regions of HS tunnels in our samples, and were also detected in sebaceous compartments in healthy skin (Figure 2, Supplementary Figure 2a).

**Figure 2.**
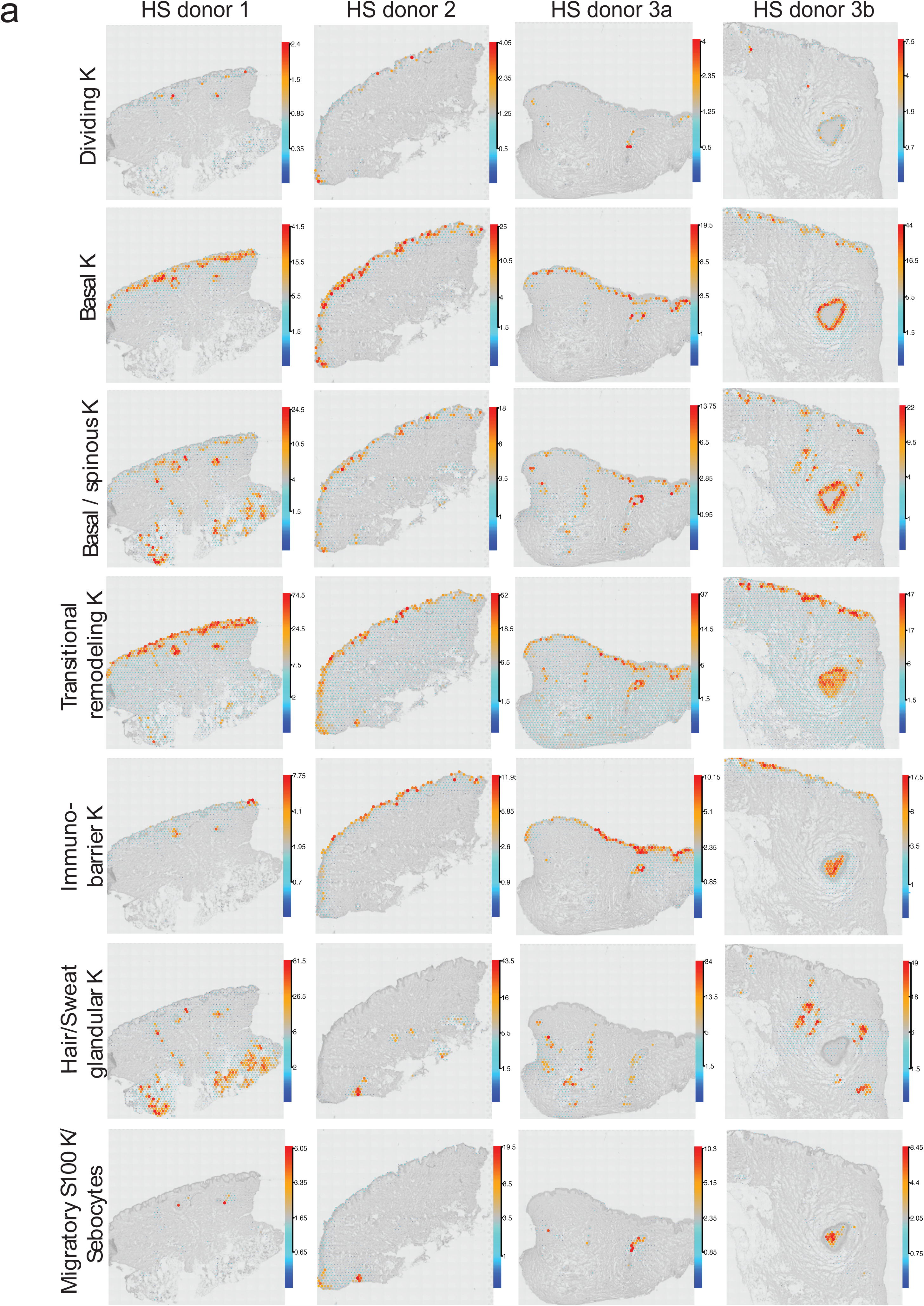
Spatial architecture of keratinocyte states in severe HS. **(a)** Spatial deconvolution using cell2location pipeline of annotated scRNAseq keratinocyte states onto Visium slides, showing localisation of 7 keratinocyte states in severe HS tissue. Predicted cell abundances shown by color gradients per spot in tissue architecture images (H&E in gray).

### Fibroblast–Keratinocyte Signalling Networks Are Striking in Severe HS Lesions

Next, we performed cell-cell interaction analysis on scRNAseq data using CellChat, which infers potential intercellular communication networks by modelling the expression of known ligand-receptor pairs between cell populations, to explore interactions associated with HS keratinocyte states. CellChat analysis identified fibroblast-keratinocyte interactions as among the strongest (Figure 3a) and most numerous (Figure 3b) predicted interactions within HS lesions. In particular, SFRP4+ reticular fibroblasts and COL6A5+ papillary fibroblasts had the highest outgoing communication score, while transitional remodeling keratinocytes, basal keratinocytes and migratory S100+ keratinocytes demonstrated the highest incoming interaction strength (Figure 3c). Top predicted signalling pathways in HS lesions included extracellular matrix-related interactions such as collagen, laminin and desmosomal pathways (Figure 3d).

**Figure 3.**
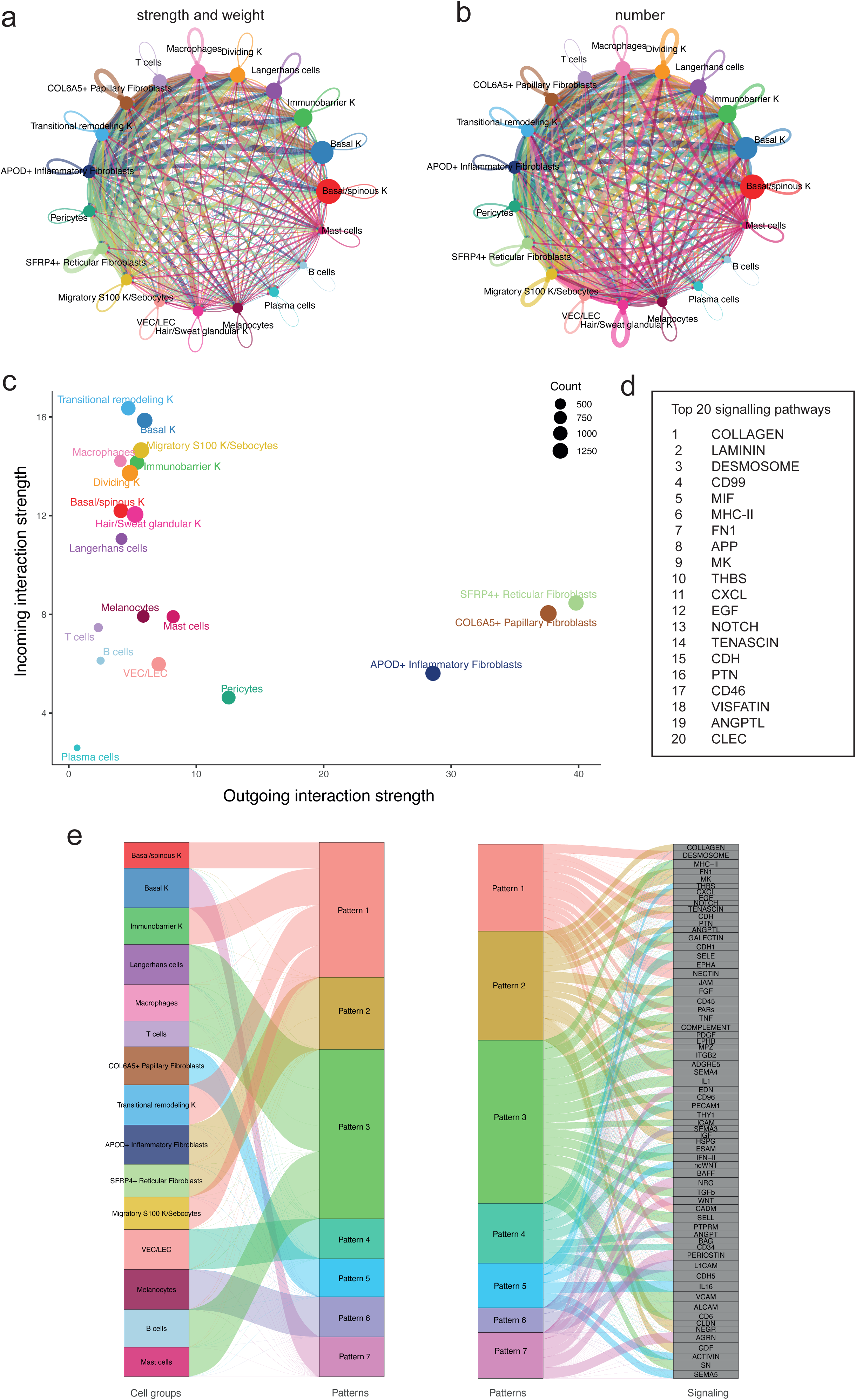
Cell–cell interactions analysis suggests fibroblast-associated signalling networks linked to pathogenic keratinocyte states in severe HS. **(a)** Circle plot of inferred intercellular communication probabilities across significant ligand–receptor pairs between cell populations in HS scRNAseq data. Each circle (node) represents a different cell population, with size indicating relative abundance. Lines (edges) connect the cell populations, illustrating communication between them, with the width or thickness indicating the strength and weight of the signaling interaction. **(b)** Circle plot showing the total number of ligand–receptor pairs between each cell population in HS scRNAseq data. **(c)** Scatter plot showing the dominant senders (sources) and receivers (targets); x-axis and y-axis are respectively the total outgoing or incoming communication probability associated with each cell population. Dot size is proportional to the number of inferred links (both outgoing and incoming) associated with each cell population. Dot colors indicate different cell populations. **(d)** Table showing the top 20 signalling path-ways in identified by CellChat in HS scRNAseq data. **(e)** River plot demonstrating inferred outgoing communication patterns of secreting cells, and the signaling pathways associated with those communication patterns.

Communication patterns analysis using CellChat, based on non-negative matrix factorisation to group coordinated ligand-receptor signalling networks, identified two distinct signalling patterns associated with fibroblast populations (Figure 3e). COL6A5+ papillary fibroblasts displayed a unique communication pattern (numbered here pattern 5), enriched for pathways such as non-canonical Wnt signalling and activin, which is part of the TGF-β superfamily. These pathways are implicated in processes such as epithelial migration, invasion and epithelial-mesenchymal transition (EMT) (Bashir et al., 2015; Liu et al., 2022; Weiss & Attisano, 2013). Recent genome-wide association and transcriptomic analyses have also implicated Wnt signaling and epithelial remodeling pathways in HS pathogenesis (Atlas Khan et al., 2025; Spindler et al., 2026). In contrast, SFRP4+ and APOD+ fibroblasts shared a second signaling pattern (pattern 2), enriched for extracellular matrix and growth signalling pathways including collagen, fibronectin, FGF and IGF.

To further define the spatial organisation of fibroblast populations within HS lesions, we performed cell2location-based projection of COL6A5+ papillary, APOD+ inflammatory and SFRP4+ reticular fibroblasts onto Visium sections across multiple HS donors (Figure 4a, 4b, 4c). These analyses demonstrated distinct spatial enrichment patterns, with COL6A5+ papillary fibroblasts preferentially localised to regions adjacent to epidermis and epithelial structures, whereas APOD+ inflammatory fibroblasts were distributed throughout the dermis. SFRP4+ reticular fibroblasts occupied deeper dermal areas, as expected. To complement this analysis, we calculated Pearson correlation coefficients (PCC) of cell-type abundances derived from the cell2location predictions across all ST samples to assess co-localisation of fibroblast populations in HS lesional skin(Figure 4d, 4e, 4f). COL6A5+ papillary fibroblasts showed the strongest spatial correlation with undifferentiated HS keratinocytes, particularly basal and dividing keratinocyte populations. In contrast, APOD+ inflammatory fibroblasts only co-localised with hair and sweat gland-associated keratinocyte populations. SFRP4+ reticular fibroblasts did not show strong co-localisation with keratinocytes, as expected for a reticular-like signature, but instead did co-localise with immune cell populations such as macrophages and mast cells (Figure 4f). Together, these findings support a prominent role for COL6A5+ papillary fibroblasts within the fibroblast–keratinocyte communication network in HS.

**Figure 4.**
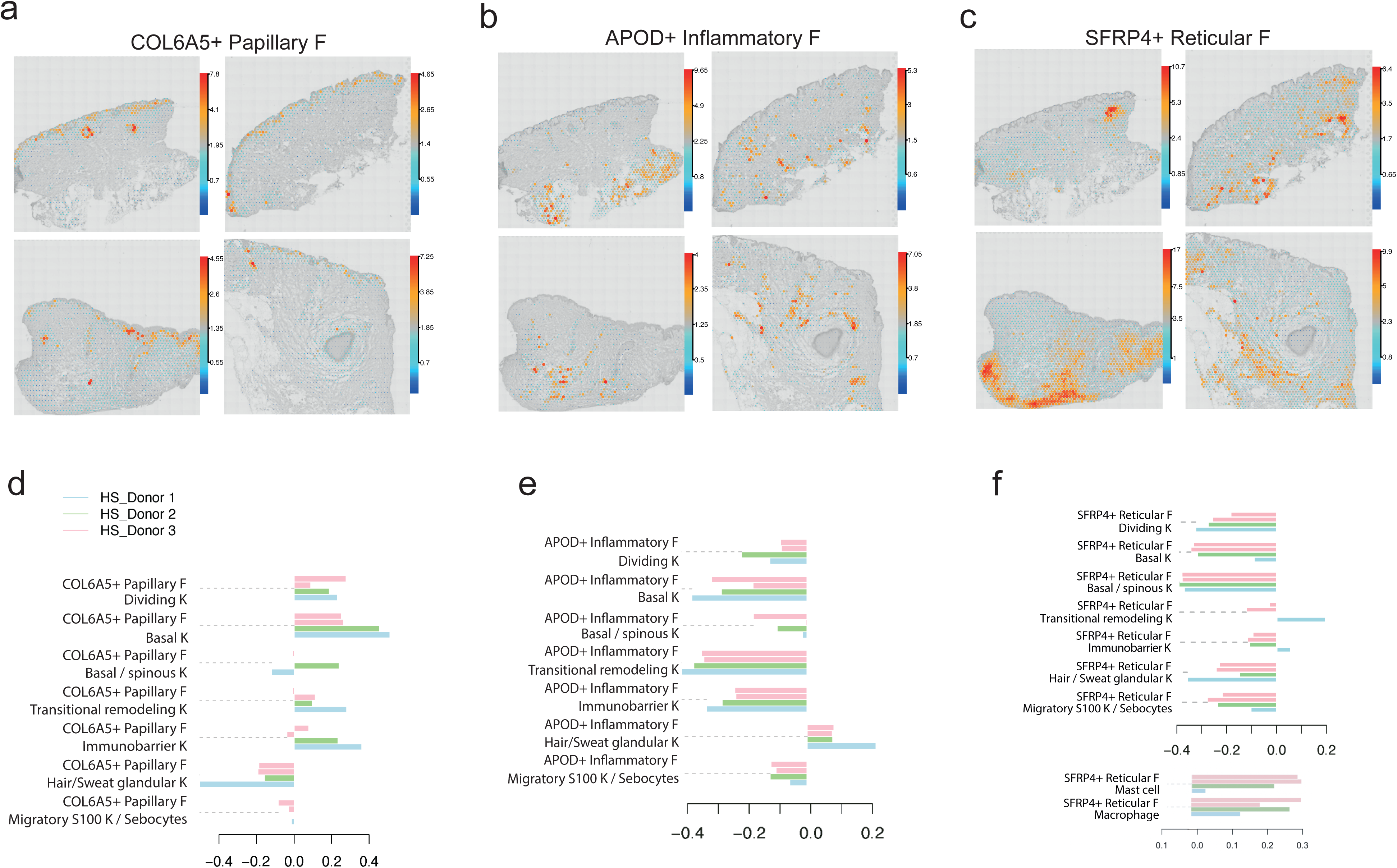
Spatial mapping reveals distinct fibroblast-associated microenvironments in HS skin. **(a-c)** Spatial projection of fibroblast subpopulations using cell2location across Visium sections from HS donors. **(a)** COL6A5+ papillary fibroblasts, **(b)** APOD+ inflammatory fibroblasts, and **(c)** SFRP4+ reticular fibroblasts show distinct spatial enrichment patterns within the dermis. **(d–f)** Spatial cell cluster co-localisation analysis using Pearson correlation coefficient (PCC) of spot-normalized cell abundances derived from cell2location predictions across Visium samples. Each bar represents an individual HS donor. **(d)** Co-localisation of COL6A5+ papillary fibroblasts with keratinocyte populations. **(e)** Co-localisation of APOD+ inflammatory fibroblasts with keratinocyte populations. **(f)** Co-localisation of SFRP4+ reticular fibroblasts with immune cell populations, including mast cells and macrophages.

### APOD+ fibroblasts are spatially associated with TLO-like immune niches in HS lesions

TLOs have recently been proposed to play an important role in HS pathogenesis (Gudjonsson et al., 2020; Lee et al., 2026; Lowe et al., 2024; Yu et al., 2024). A key feature of TLOs is clonal expansion of plasma cells and proliferating B cells. In our analysis, plasma/B cells were relatively abundant and are over-represented in HS compared to other inflammatory skin diseases (Supplementary Figure 2b). Further analysis of these populations identified an activated B cell phenotype with increased expression of AICDA, amongst other genes (Figure 5a-b) (Horns et al., 2020). AICDA is responsible for somatic hypermutation and class-switch recombination, processes that typically facilitate antibody diversification in secondary lymphoid tissues (Honjo et al., 2002). To further characterise the spatial organisation of immune cells in HS lesions, we performed immunofluorescence staining for lymphocyte and antigen-presenting cell markers (Figure 5d-f). This demonstrated the presence of lymphocyte-rich aggregates within HS dermis, containing B cells, T cells and antigen-presenting cells, with expression of activation and plasma-associated markers. These findings support the presence of organised immune aggregates with features consistent with TLO-like structures. . Plasma and B cells were mapped onto Visium sections using cell2location (Figure 5c) and plotting of plasma cell markers (IGHA1, IGHM, IGHG1) showed enrichment of plasma / B cells surrounding HS lesional structures, with high expression of IgA and IgG immunoglobulins in ST (Supplementary Figure 2d), consistent with the presence of IgA and IgG-expressing plasma cells in scRNAseq (Supplementary Figure 2e), although relative abundance varied between samples

**Figure 5.**
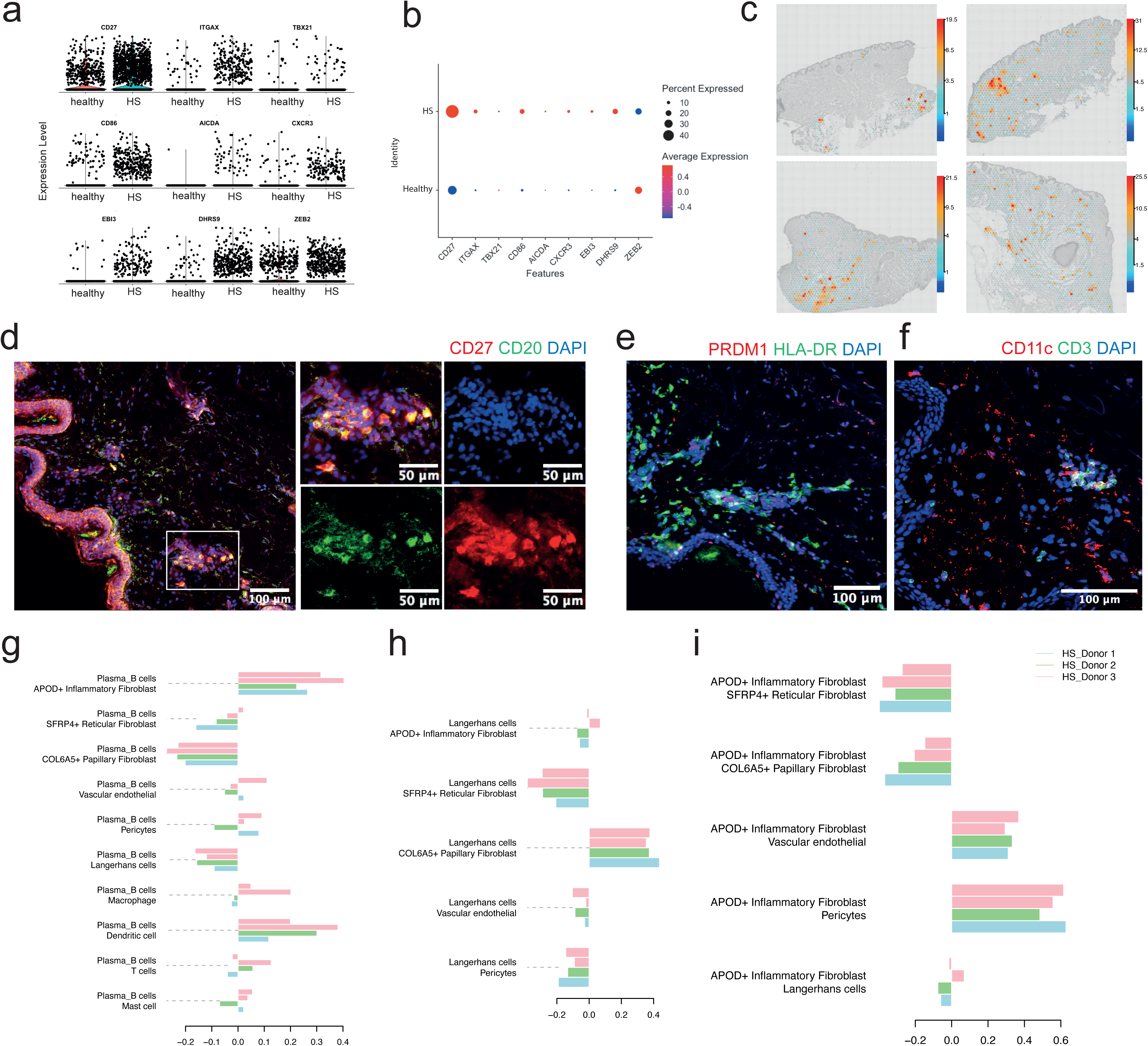
Activated B and plasma cells localise with immune aggregates associated with fibroblasts and immune microenvironments in HS skin, consistent with tertiary lymphoid-like organisation in HS. **(a)** Violin plots of gene expression of B cell markers from HS scRNAseq compared to publicly available healthy skin scRNAseq dataset (Ganier et al. 2024) representing an activated B cell phenotype in HS. Each dot is a cell, and the height of dot on the y axis is the normalized expression level. **(b)** Dot plot showing expression of B cell markers in HS versus healthy skin (Ganier et al. 2024) confirming this activated B cell phenotype in HS. **(c)** Spatial deconvolution using cell2location of annotated scRNAseq plasma/B cells onto Visium slides, confirmed their localisation in severe HS tissue. Predicted cell abundanc-es shown by color gradients per spot in tissue architecture images (H&E in gray). **(d)** Immunofluorescent staining of HS tissue showing dermal aggregates enriched for CD20+ B cells (green) and CD27+ activated/memory B cells (red). Low-magnification overview (left) with higher-magnification inset (right) demonstrates clustering of B cells wtihin inflammatory aggregates. Split-channel images confirm the spatial distribution of CD20 and CD27 signals wtihin these areas. **(e)** Immunofluorescent staining of HS tissue showing PRDM1+ cells (red) and HLA-DR+ antigen-presenting cells (green) within dermal inflammatory aggregates. These regions demonstrate the presence of plasma cell-associated nad antigen-presenting cell populations within the same spatial compartments. **(f)** Immunofluorescent staining showing CD11c+ dendritic cells (red) and CD3+ T cells (green) within dermal inflammatory aggregates, indicating the presence of mixed immune cell populations within these structures. **(g)** Spatial cell cluster colocalization analysis for plasma/B cells with all fibroblast populations, immune cells and endothelial cells using PCC analysis. The bar chart shows PCC of spot normalized cell abundances using the spatial cell2location predictions across all spots of Visium samples, with each individual bar representing a Visium sample (HS donor). **(h)** Spatial cell cluster colocalization analysis for APOD+ fibroblasts with other fibroblast subpopulations, endothelial cells, pericytes and langerhans cells using PCC analysis. **(i)** Spatial cell cluster colocalization analysis for Langerhans cells with fibroblast subpopulations, endothelial cells and pericytes using PCC analysis.

We utilised PCC to assess the co-localisation of cell populations identified by cell2location. Plasma and B cells showed strongest colocalisation with APOD+ fibroblasts and dendritic cells, and to a lesser extent mast cells and T cells (Figure 5g). Further analysis showed that APOD+ inflammatory fibroblasts also co-localised with vascular endothelial cells and pericytes (Figure 5h). In scRNAseq data, endothelial cells in HS were found to have high levels of ACKR1, a marker of venular endothelial cells which encodes a membrane protein that facilitates immune cell trafficking and is also observed in other inflammatory skin diseases (Reynolds et al., 2021). In lesional HS, the expression of ACKR1 expression was proportionally greater than that observed in endothelial cells in lesional eczema and psoriasis (Supplementary Figure 2c). Together, these findings are consistent with the presence of organised immune niches in HS lesional tissue (Figure 5d-f) and suggest that APOD+ inflammatory fibroblasts are spatially associated with these aggregates, in keeping with recently proposed TLO-like structures in HS.

Next, we explored the signaling interactions within these immune niches using CellChat. CellChat analysis showed that Langerhans cells, T cells, dendritic cells, macrophages, B cells and mast cells were associated with the most expansive communication pattern (Pattern 3) which includes canonical HS cytokine signatures such as TNF, IL-1 and TGF-β (Figure 3e). Plasma/B cells did not show strong interaction strength either as senders or receivers (Figure 3c) in scRNAseq cell-cell interaction analysis, and plasma cells were not prominent in communication pattern analysis (Figure 3e). This is consistent with a model in which plasma cells may contribute to HS inflammation indirectly through the production of antibodies, rather than through direct signalling to keratinocytes.

Recently, Langerhans cells have been identified as being important coordinators of TLO formation in mouse skin, in response to colonising micro-organisms (Gribonika et al., 2025). In our data, Langerhans cells showed the highest incoming interaction score among immune populations (Figure 3c). Furthermore, Langerhans cells were predicted to co-localise with COL6A5+ papillary fibroblasts as well as pathogenic HS keratinocytes (Figure 5i). This suggests that Langerhans cells may act as an interface between keratinocytes populations forming pathogenic HS lesions, and immune niches resembling TLO-like structures, potentially linking epithelial remodelling with sustained inflammation.

Overall, our findings suggest that presence of two distinct niches in HS lesions: COL6A5+ fibroblasts, which are associated with pathogenic keratinocyte states, and APOD+ fibroblasts, which localise with immune cell aggregates and vascular structures in keeping with a role in supporting chronic inflammatory niches.

## Discussion

The genetic architecture of HS consistently implicates epidermal differentiation in its pathophysiology (Kjærsgaard Andersen et al., 2025). Recent work by Navrazhina et al. and Jin et al. has placed keratinocytes at the centre of HS pathogenesis (L. Jin et al., 2023; Navrazhina et al., 2021). Navrazhina et al. identified tunnel epithelium as being chemotactic and pro-inflammatory, whilst Jin et al. identified a disease-specific keratinocyte state arising from epigenetic reprogramming of basal keratinocytes towards a hyperproliferative and pro-inflammatory phenotype in HS (L. Jin et al., 2023).

Although pro-fibrotic and pro-inflammatory HS fibroblast populations have previously been described in HS (van Straalen et al., 2024), the spatial relationships and signalling interactions between fibroblast populations and pathogenic keratinocytes have not been well defined. In HS, keratinocytes breach their physiological compartment and invade the dermis, forming 3-dimensional structures representing inflammatory epithelial niches. In our data, COL6A5+ papillary fibroblasts co-localised with basal keratinocytes, and showed strong predicted signalling interactions with these populations, consistent with a role in supporting the migratory and proliferative pathogenic keratinocyte states associated with HS lesions. This interaction is reminiscent of developmental epithelial-mesenchymal signalling seen in skin appendage formation, where papillary fibroblasts are required for the formation of structures such as pilosebaceous units (Driskell et al., 2013; Philippeos et al., 2018). These observations suggest that the formation of epithelial structures seen in severe HS may require coordinated remodelling of the surrounding connective tissue to permit their physical extension into the dermis. We also identified a second fibroblast niche characterised by APOD+ fibroblasts, which co-localised with endothelial cells, pericytes, plasma cells and B cells. This pattern is consistent with the organisation of TLO-like structures, which are increasingly recognised in HS and may contribute to the maintenance of chronic inflammation in the HS tissue microenvironment. Recent spatial transcriptomic analysis of early HS lesions has shown that B-cell and T-cell activation is already present prior to the development of tunnels and fibrosis, suggesting that immune niche formation may occur early and subsequently expand during disease progression (Lee et al., 2026).

Abnormal keratinisation and barrier alteration are recognised as key factors in the development of HS (Frew, 2020; Zouboulis et al., 2020). Barrier integrity influences the composition of the skin microbiome and the activation of innate immune responses (Belkaid & Tamoutounour, 2016). The role of the microbiome in HS pathogenesis is supported by the efficacy of antibiotics in suppressing HS inflammation and their continued use as first-line HS therapy (Sabat et al., 2020), as well as by studies showing that microbial shifts are detectable in non-lesional HS tissue and become more pronounced as disease progresses (Ring et al., 2017). Recent work has also demonstrated that gram-negative anaerobes enriched in HS lesions can directly activate keratinocytes (Agbogan et al., 2026; Williams et al., 2024), supporting the hypothesis that microbial signals may precede clinically apparent disease. In this context, it is notable that Langerhans cells, which survey the skin barrier for harmful microbes, are localised within the COL6A5+ fibroblast niche associated with early keratinocyte remodeling in our data. Langerhans cells have been shown to express genes related to the NLRP3 inflammasome (Moran et al., 2023), a pathway strongly implicated in HS pathogenesis (Johnston et al., 2021) and have recently been identified as coordinators of bacteria-induced TLO structures in skin (Gribonika et al., 2025). Together, these observations raise the possibility that Langerhans cells may represent an interface between epithelial remodelling and barrier-associated immune activation, linking early keratinocyte changes with the development of chronic inflammatory niches in HS. Further functional studies are required to determine whether Langerhans cells contribute to disease initiation, persistence, or both.

Current therapeutic strategies in HS primarily target immune pathways, including TNF-alpha and IL-17. However, treatment responses remain dissatisfactory for patients (Garg et al., 2020) and primary or secondary failures are seen relatively frequently in HS (Aarts et al., 2021). The expanding therapeutic pipeline, including agents targeting JAK, BTK and IL-1 pathways, further reflects the biological heterogenity of HS and the likelihood that multiple pathogenic mechanisms contribute to disease activity. Our data support the presence of two distinct fibroblast niches within the HS tissue microenvironment, reflecting separate cellular networks associated with epithelial remodeling and chronic immune activation in HS pathogenesis. These findings suggest that targeting individual inflammatory pathways alone may be insufficient to fully control disease activity or modify disease progression, as HS pathogenesis appears to involve coordinated epithelial, stromal and immune interactions within the lesional microenvironment. Future therapies may therefore need to address both early epithelial dysfunction and stromal-immune niches that establish the HS tissue microenvironment in order to alter the natural history of the disease.

### Ethics statement

The study was sponsored by Guy’s and St Thomas’ NHS Foundation Trust and King’s College London and was subject to both institutional and external research ethics council (REC) review (REC reference 19/NE/0063).

## Materials and Methods

### Patients and samples

Lesional HS skin from 3 donors was obtained from wide local excision surgery following ethical approval (REC reference 19/NE/0063). Patients provided written informed consent prior to surgery.

### Generation of HS single-cell RNA-seq and spatial transcriptomic data

HS skin was enzymatically dissociated to separate epidermis and dermis, followed by preparation of single-cell suspensions using standard procedures. Cells were stained with a viability dye, sorted by FACS, and assessed for viability prior to loading on the 10x Genomics Chromium platform (Single Cell 3′). Libraries were sequenced on an Illumina HiSeq 4000 and processed with CellRanger against GRCh38.

Fresh-frozen HS skin was embedded in OCT, cryo-sectioned, and processed on 10x Genomics Visium (Spatial 3′). RNA integrity was assessed using RNAscope with housekeeping control probes. Library quality was verified prior to sequencing on an Illumina HiSeq 4000. Reads were processed using spaceranger with GRCh38.

Further information regarding digestion conditions, instrument models, probe sets, fluorophores, and tissue optimization parameters are detailed in the *Supplementary Materials and Methods*.

### ScRNAseq and spatial transcriptomic data analysis

scRNA-seq and spatial data underwent quality control, normalization, dimensionality reduction, and clustering in Seurat v5.1 (Hao et al., 2024). Cell types were assigned by canonical marker genes and gene ontology of cluster-enriched features. Datasets were integrated using Harmony (Korsunsky et al., 2019) for downstream analyses.

Cell type abundances across Visium spots were inferred using cell2location trained on the scRNA-seq reference. To identify tissue microenvironments characterized by co-occurring cell types, we applied non-negative matrix factorization (NMF) to normalized cell type abundance estimates across Visium spots. Further details regarding cell2location and colocalisation analysis are described in the *Supplementary Materials and Methods*.

Keratinocyte lineage dynamics were analyzed with Monocle 3 (Cao et al., 2019). Cells were converted from Seurat to Monocle, partitioned, and ordered in pseudotime with the basal keratinocyte population set as the root state. Ligand–receptor-mediated signaling between cell states was inferred using CellChat (S. Jin et al., 2025) on the annotated scRNA-seq dataset. We computed communication probabilities and performed pathway-level input/output analyses using the standard pipeline.

### Immunohistochemistry and microscopy

Skin samples were embedded in OCT and cryo-sectioned at 10–16 µm thickness. Sections were fixed in 4% paraformaldehyde, blocked in serum-based buffer, and incubated with primary antibodies overnight at 4 °C. After washing, sections were labeled with secondary antibodies and DAPI, then mounted with antifade medium. Full reagent details, concentrations, incubation times, and microscopy analysis details are provided in *Supplementary Materials and Methods*.

### Data Availability Statement

The raw and processed datasets for scRNAseq and Visium ST generated in the current study are available on ArrayExpress.

### Conflict of Interest Statement

XDH has received honoraria from AbbVie; served in advisory roles for RoC Skincare, Dermatica Ltd; and undertaken advisory and promotional roles for Simple Skincare, E45, Skin + Me Ltd., L’Oreal Dermatological Beauty and The Skin Diary Ltd. ER has served on advisory boards for Biogen, Novartis, and UCB; as investigator for trials sponsored by Boehringer Ingelheim, Novartis, Incyte, UCB, and Abbvie; received research funding from UCB; serves as medical director of Klira Ltd; and has undertaken promotional work for CeraVe, La Roche Posay, Eucerin, Paula’s Choice, E45, Roc, and Garnier. MDL and FMW are co-founders of Fibrodyne Ltd, a company working on fibroblast cell therapies.

## Acknowledgements

XDH was the recipient of an Accelerator Award from Cancer Research United Kingdom, which supported this work. FMW acknowledges financial support from the United Kingdom Medical Research Council (MR/ PO18823/1), Biotechnology and Biological Sciences Research Council (BB/M007219/1), and the Wellcome Trust (206439/Z/17/Z) at the time of this work. This work was supported by the Francis Crick Institute, which receives its core funding from Cancer Research United Kingdom (FC010110), the United Kingdom Medical Research Council Journal of Investigative Dermatology (FC010110), and the Wellcome Trust (FC010110). During this project, NML was a Winton Group leader in recognition of the Winton Charitable Foundation’s support toward the establishment of the Francis Crick Institute. NML was also funded by a Wellcome Trust Joint Investigator Award (103760/Z/14/Z), the Medical Research Council eMedLab Medical Bioinformatics Infrastructure Award (MR/L016311/1), and received core funding from the Okinawa Institute of Science and Technology Graduate University. MDL gratefully acknowledges financial support from the Wellcome Trust (211276/E/18/Z).

The Advanced Sequencing Facility at the Francis Crick Institute provided protocols, technical support and guidance for single-cell RNA sequencing experiments. The Genome Centre, Queen Mary, University of London provided support with their genomics platform for spatial transcriptomics. Support for confocal microscopy was provided by King’s College London Nikon Centre and Dr Ines Tomas.

## Author Contribution Statement

**XDH:** Conceptualization, Data Curation, Formal Analysis, Funding Acquisition, Investigation, Methodology, Project Administration, Visualization, Writing – original draft, Writing – review and editing

**CG:** Conceptualization, Formal Analysis, Investigation, Methodology, Project Administration, Visualization, Supervision, Writing – original draft, Writing – review and editing

**PM:** Data Curation, Software, Resources

**CC:** Data Curation, Resources

**JG:** Investigation

**ER:** Resources

**NML:** Conceptualization, Resources, Supervision, Writing – review & editing

**MDL:** Conceptualization, Methodology, Supervision, Visualization, Writing – review & editing

**FMW:** Conceptualization, Methodology, Resources, Supervision, Visualization, Writing – original draft, Writing – review & editing

## Notes

### Summary of Updates

In this revised version, we have updated the manuscript to improve clarity and interpretation in response to reviewer feedback. We have moderated language to better reflect that conclusions are based on spatial association and inferred interactions rather than direct functional evidence. We have added immunofluorescence experiments to provide orthogonal support for the presence and organisation of immune cell aggregates within HS tissue. We have also improved the presentation and description of spatial analyses, particularly for fibroblast populations, and updated figure layout and legends accordingly. Finally, we have expanded the Discussion to better contextualise our findings within recent single-cell studies of HS and corrected minor errors in figure labelling and text.

