## Supplemental Material and Methods for "Pathogenic Keratinocyte States and Fibroblast Niches Define the Tissue Microenvironment in Severe Hidradenitis Suppurativa"

### Supplemental Materials and Methods

#### Generation of HS single-cell RNA-seq and spatial transcriptomic data

##### Generation of single-cell RNA-seq data:

Skin was enzymatically digested with Dispase overnight at 4°C to separate epidermis from dermis. Subsequently, the epidermis was digested with a mixture of trypsin-EDTA and Versene for 15 minutes at 37^o^C. The dermis was dissociated using the Whole Skin Dissociation kit for human material (Miltenyi Biotec, cat. no. 130-101-540) through incubation with the enzyme solution for 3 hours in a shaking water bath at 37°C. Fetal bovine serum (FBS) was then added to halt the digestion, and the cell suspensions were filtered through 70-µm and 40µm Falcon cell strainers. Cell suspensions were centrifuged at 1200 rpm for 4 minutes at 4°C and the supernatant was aspirated. The collected cells were frozen in 10% DMSO in FBS and stored in liquid nitrogen tanks. For library preparation, cells were thawed and resuspended in PBS with 2% FBS.

Receiving 1.5ml Eppendorf tubes containing 50µl 1X PBS with 2% FBS were placed on ice. Single cell suspensions were stained with 1µl 0.05µl/ml DAPI prior to sorting. Cell sorting was performed on the BD FACSAria II and BD FACSAria III Fusion cell sorters in the King’s College London Biomedical Research Centre (BRC). After sorting, the suspension was spun down at 4^o^C at 1200 rpm for 4 mins, and supernatant was aspirated to leave approximately 70µl of suspension for quality control and scRNAseq library preparation.

Suspension quality was assessed by staining a small sample with Trypan blue and loading this onto the EVE cell counter (NanoEnTek). After passing initial QC with a 70% minimum viable cell cut-off, the suspension was loaded onto a Chromium chip and underwent droplet encapsulation on the Chromium controller from 10X Genomics (Pleasanton CA, USA). The sequencing of the libraries was carried out by staff at the Advanced Sequencing Facility of the Francis Crick Institute using Single Cell 3’ reagent kits following the manufacturer’s protocol. Library quality was assessed using Agilent 4200 Tapestation prior to sequencing. Finally, libraries were sequenced using an Illumina HiSeq 4000 device. Raw scRNAseq data was run through CellRanger, v6.1.1 (10X Genomics) using the GRCh38 reference genome.

##### Generation of spatial transcriptomic data

Optimal RNA integrity of the skin sections was assessed using the RNAscope Multiplex Fluorescent Detection Kit v2 (cat. no. 323100; ACDBio, Newark, California), following the manufacturer’s instructions. Skin sections were assessed with RNAscope housekeeping control probes (high (UBC), medium (PPIB) and low (POLR2A) expressors). Probes against targeted human mRNA molecules were used (all from ACDBio catalog probes). Opal dyes (Akoya Biosciences, Marlborough, Massachusetts) were used at a dilution of 1:1,000 for the fluorophore step to develop each channel: Opal 520 Reagent Pack (FP1487001KT), Opal 570 Reagent Pack (FP1488001KT) and Opal 650 Reagent Pack (FP1496001KT). Nuclei were counterstained with 4′,6-diamidino-2-phenylindole and mounted using ProLong Gold Antifade Mountant (ThermoFisher, Canoga Park, California, cat. no. P36930). Fresh frozen skin samples were embedded in OCT, cryo-sectioned and processed according to the recommended protocols (Tissue optimization: CG000238; Gene expression: CG000239) using the Spatial 3’ v1 kit. Optimal permeabilisation time for 10um skin sections was determined to be 20 minutes. cDNA libraries were quality controlled using the Agilent Bioanalyser prior to sequencing on the Illumina HiSeq 4000 system. Raw data was run through spaceranger-1.3.1, with alignment performed against the GRCh38 human reference genome.

#### ScRNAseq and spatial transcriptomic data analysis

##### Deconvolution of scRNAseq clusters into 10X Visium sections using cell2location

To map scRNA-seq clusters onto 10X Visium spatial transcriptomics sections, we employed cell2location v0.1 (Kleshchevnikov et al. 2022). This process begins with training a negative binomial regression model to infer reference transcriptomic signatures for each cell type identified in the scRNA-seq dataset. These reference profiles are then used to estimate cell type abundances across the spatial transcriptomics (ST) slides. Prior to model training, genes with very low expression levels were excluded. The training was run for 50,000 iterations. The following cell2location hyperparameters were used: (1) expected cell abundance (N_cells_per_location) = 30; (2) regularisation strength of detection efficiency effect (detection_alpha) = 20. All other parameters were used at default settings.

##### Co-localisation analysis

To detect tissue microenvironments characterized by co-localized cell types, we applied non-negative matrix factorization (NMF). Initially, we normalized the estimated cell type abundance matrix by dividing each value by the total abundance per spot. This produced a normalized matrix Xn with dimensions 𝑛 × 𝑐, where n represents the number of spatial spots across Visium slides and c denotes the number of reference cell types. We then decomposed this matrix using the formula 𝑋n = 𝑊𝑍, where W is an 𝑛 × 𝑑 matrix containing latent factor values for each spot, and Z is a 𝑑 × 𝑐 matrix indicating the proportion of each cell type's abundance associated with each latent factor. These latent factors reflect distinct tissue microenvironments defined by specific combinations of cell types.

The decomposition was performed using the NMF package in R (Gaujoux and Seoighe 2010), with the number of factors 𝑑 set to Hi NN and the default algorithm employed. To standardize the NMF coefficients, we normalized them by the maximum value within each factor. We executed the NMF process 100 times and generated a coincidence matrix to assess consistency across runs. The optimal run was selected based on the lowest mean silhouette score calculated from the coincidence matrix. In cases where multiple runs shared the minimum silhouette score, the one with the lowest deviance (as reported by the nmf function) was chosen. For analyzing correlations in cell type abundance, we used the normalized Xn matrix. Pearson correlation coefficients (PCCs) were computed for every pair of cell types within each sample

#### Immunohistochemistry and microscopy

##### Immunohistochemistry

Skin tissue samples were embedded in Tissue-Tek O.C.T. (Life Technologies, Waltham, MA) and stored at –80°C prior sectioning. Sections of 10-16 micron were cut using a Thermo Cryostar Nx70 (Thermo Fisher Scientific, Waltham, MA). Sections were stained with Harris’ hematoxylin solution for 20s at room temperature and were then rinsed in tap water. Next, 0.225% Acid Alcohol (acetic acid and ethanol) in water was used to differentiate the tissue for 10s followed by rinsing with tap water. In the bluing step, tissue was soaked with Scott’s tap water (ref: 3802900, Leica Biosystems, Milton keynes, UK) for 10s and then rinsed with tap water. Staining was performed with 0.125% eosin Y ethanol solution for 10s. Finally, skin sections were dehydrated in IMS for 20s, then cleared in Xylene 30s and mounted with CV mount medium (ref: 14046430011, Leica biosystem, Germany). For immunohistochemistry, sections were fixed in 4% paraformaldehyde, blocked with a solution of 10% donkey serum, 0.1% fish skin gelatin, 0.1% Triton X-100, and 0.5% Tween20 (all from Sigma-Aldrich, St. Louis, MO) in 1x phosphate buffered saline (PBS) and labelled with primary antibodies diluted in blocking buffer overnight at 4°C. Sections were then washed with 1x PBS and labelled with secondary antibodies and DAPI for 1 hour at room temperature, washed with 1x PBS, and mounted with ProLong™ Gold Antifade Mountant (ThermoFisher).

##### Microscopy imaging and analysis

Confocal microscopy was performed with a Nikon A1 Upright Confocal microscope (Tokyo, Japan) using 10x or 20x objectives at the Nikon Centre, King’s College London. Image processing was performed with Image J (Fiji) (National Institutes of Health, Bethesda, MD). Imaging of hematoxylin and eosin-stained sections was performed using a Hamamatsu NanoZoomer slide scanner (Hamamatsu City, Japan) and image processing was performed with NDP.view2 (Hamamatsu City, Japan).
