## Supplementary Figure 1 for "Pathogenic Keratinocyte States and Fibroblast Niches Define the Tissue Microenvironment in Severe Hidradenitis Suppurativa"

a

| Patient | Relevant medical history | Age at donation | Sex | FH | Affected sites | Age of onset | Smoking history |
|---|---|---|---|---|---|---|---|
| HS1 | Hypertension | 54 | Female | Yes | Axillae and sometimes groin | 20s | Yes |
| HS2 | Thalassaemia trait | 22 | Female | Yes | All flexural sites | Teens | No |
| HS3 | Previous gastric bypass | 52 | Female | Nil | All flexural sites | 40s | Yes |

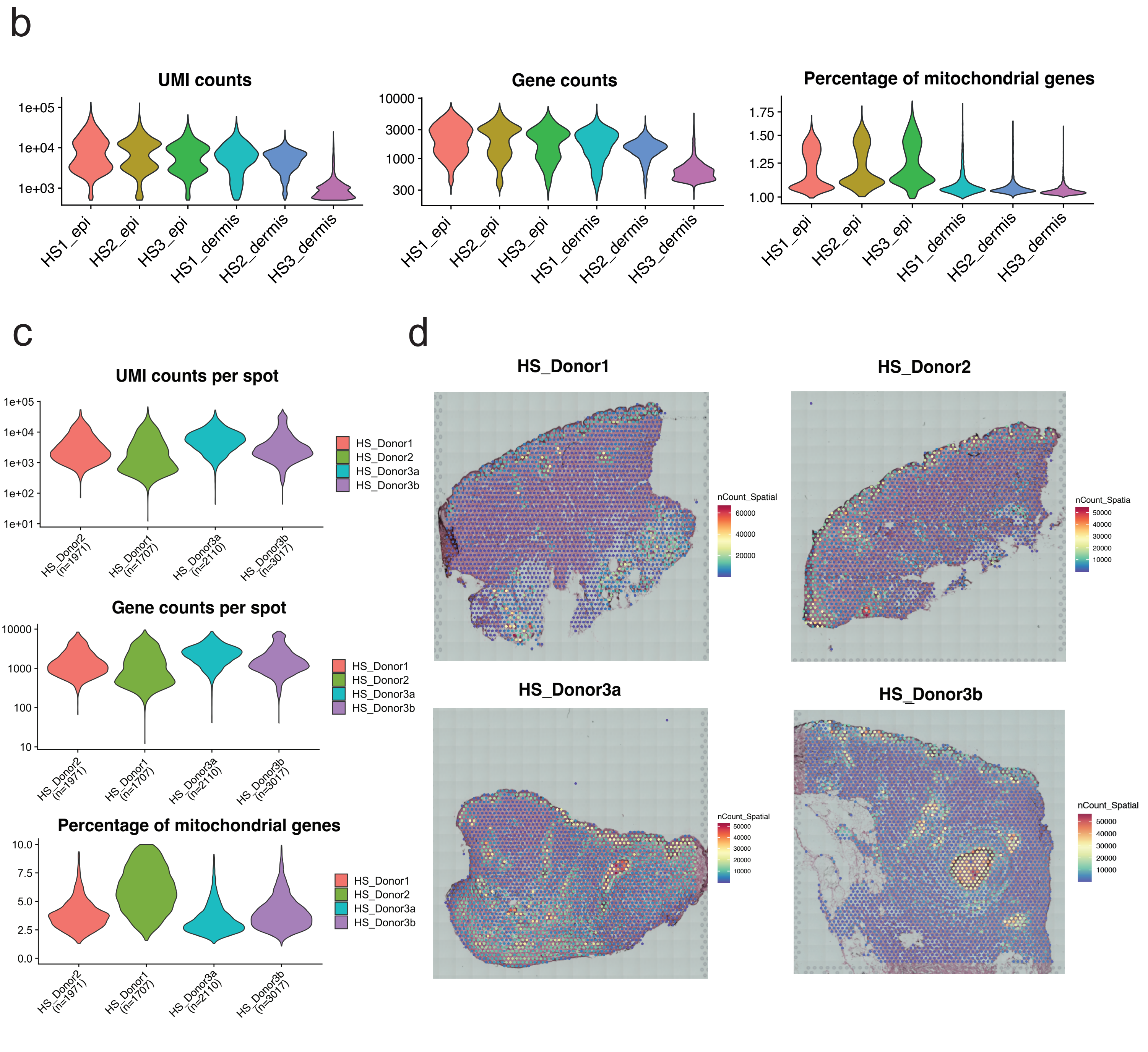


**Supplementary Figure 1. Metadata and quality control metrics of scRNAseq and spatial transcriptomics in HS samples. (a)** Relevant metadata from 3 donors with severe HS. **(b)** Violin plots of key quality control parameters related to scRNAseq data from 3 donors with severe HS, showing frequency distribution of log1p-transformed unique molecular identifier (UMI) counts (left panel), gene counts (middle panel) and percentage of mitochondrial genes (right panel). **(c)** Violin plots of quality control parameters related to Visium data across all samples, showing frequency distribution of UMI counts (log1p-transformed) per spot (upper panel), gene counts per spot (middle panel) and percentage of mitochondrial genes per spot (lower panel). **(d)** Spatial distribution of UMI counts (nCount_spatial) across all HS tissue sections processed with the 10X Visium protocol.
