## Supplementary Figure 2 for "Pathogenic Keratinocyte States and Fibroblast Niches Define the Tissue Microenvironment in Severe Hidradenitis Suppurativa"

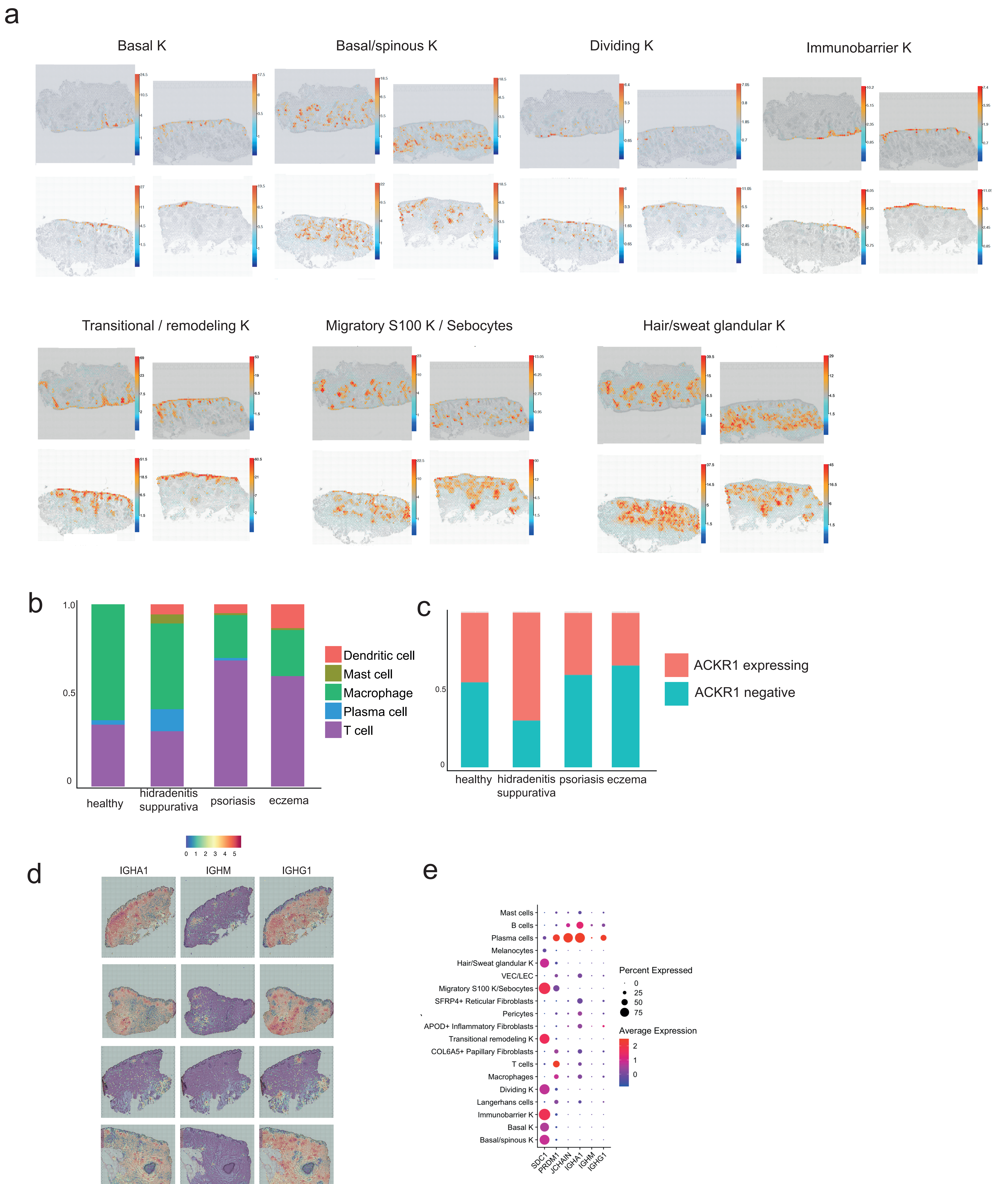


**Supplementary Figure 2. Spatial mapping of keratinocyte states in healthy human skin and comparison of immune and endothelial cellular populations in HS, Eczema, Psoriasis, and Healthy Skin. (a)** Spatial deconvolution of annotated scRNAseq keratinocyte states onto Visium slides of publicly available healthy skin scRNAseq dataset (Ganier et al. 2024) using cell2location, showing localisation of 7 keratinocyte states in healthy skin tissue. Predicted cell abundances shown by color gradients per spot in tissue architecture images (H&E in gray). Dividing K are seen scattered within basal interfollicular epidermis (IFE). Basal K are identified in the basal layer of IFE. Basal/spinous K were located in between basal and spinous layers of the IFE and throughout the pilosebaceous unit (PSU). Transitional remodeling K were deconvoluted throughout the IFE layers, except for the granular layer and extending into the upper PSU. Immunobarrier K were spatially consistent with the most differentiated and granular layer of the IFE. Hair and sweat glandular keratinocytes were seen in the lower PSU. Migratory S100+ K / sebocytes were identified within sebaceous glands of the PSU. **(b)** Analysis of proportions of cellular populations in HS scRNAseq data compared to lesional eczema, lesional psoriasis and healthy skin from (Reynolds et al. 2021). Stacked bar chart showing relative proportions of immune cell populations in scRNAseq in those datasets. T cells are proportionally higher in psoriasis and eczema skin, whereas plasma cells and mast cells are proportionally higher in HS skin. **(c)** Endothelial cells from lesional eczema skin, HS skin, psoriasis skin and healthy skin were examined for presence or absence of ACKR1 expression on a per cell basis. Proportions of ACKR1-expressing cells against ACKR1-negative cells from each dataset are shown in a stacked bar chart. **(d)** Gene expression of IGHA1, IGHM, IGHG1, representing antibody production by B/plasma cells in Visium data is detected in most HS samples throughout the dermis, with color gradient indicating gene expression level. **(e)** Dot plot of genes representing plasma cell markers (SDC1, PRDM1, JCHAIN) and antibody production (IGHA1, IGHM, IGHG1) in HS scRNAseq dataset confirming this antibody production mainly by plasma cells but also by B cells in HS skin.
